# Gut-liver-axis microphysiological system for studying cellular fluidic shear stress and inter-tissue interaction

**DOI:** 10.1101/2022.01.31.478467

**Authors:** Jiandong Yang, Yoshikazu Hirai, Satoshi Imamura, Toshiyuki Tsuchiya, Osamu Tabata, Ken-ichiro Kamei

## Abstract

Gut-liver-axis (GLA) is a fundamental interaction between the gut and liver for maintaining human health. To clarify the physiological and pathological roles of GLA in the human body, a GLA microphysiological system (GLA-MPS) holds great potential. However, in current GLA-MPS, the importance of a physiologically relevant flow for gut and liver cells’ cultivation is not fully addressed. In addition, the integration of individual organ perfusion, circulation flow, and organ tissue functions in a single device has not been achieved. Here, we introduce a GLA-MPS by integrating two cell culture chambers with individually applied perfusion flows and a circulation channel with an on-chip pneumatic micropump under cell culture chambers via a porous membrane for interconnecting them. We analyzed the fluid shear stress (FSS) with computational fluid dynamics simulations and confirmed that the physiologically relevant FSS (i.e., 8 × 10^−3^ and 1.2 × 10^−7^ dyne cm^−2^) could be applied for the gut (Caco-2) and liver (HepG2) cells, respectively. Under physiologically relevant flow, the Caco-2 and HepG2 cells in the GLA-MPS maintained a cell survival rate of 95% and 92%, respectively; further, they enhanced the expression of functional proteins such as zonula occludens 1 (ZO-1) and albumin (ALB), respectively. Thus, the presented GLA-MPS can be adapted as an advanced *in vitro* model in a wide range of applications for disease modeling associated with inter-tissue interactions, such as fatty liver diseases.

## Introduction

The gut-liver-axis (GLA) is a bidirectional interaction between the gut and liver mediated by dietary, genetic, and environmental factors,^1^ that regulates the absorption/prevention of exogenous agents at the gut barrier and metabolic processes in the liver.^2,3^ The disruption of GLA functions leads to liver disease progression and gastrointestinal disorders.^4^ Therefore, there are ongoing trends in the study of GLA for disease modeling and drug discovery. However, using experimental animal models does not yield clear information about GLA because they have a whole set of organs that show complicated interactions in addition to GLA. Therefore, it is necessary to establish an *in vitro* model to better understand GLA.

Microphysiological systems (MPSs) or organs-on-chips (OoCs) recapitulate pathophysiological conditions *in vitro* by utilizing micro/nanotechnology approaches for applications in drug discovery. The MPSs provide physiologically relevant cellular microenvironments (i.e., soluble factors, perfusion, extracellular matrices, tissue-like structures, and physical stimuli) for cells and organs,^5^ and interconnecting organs via flow channels and controls.^6^ These features are critical to investigate the absorption, distribution, metabolism, excretion, and toxicity (ADME-T) of the drug candidate and for disease modeling as well.^6^ Therefore, MPSs are suitable for recapitulating GLA *in vitro*.

Thus far, a number of MPSs as an *in vitro* GLA model (GLA-MPS) have been reported, such as GLA for metabolism and inflammation axess.^3,7– 11^ They showed better predictive capabilities of organ to organ interaction, compared to normal static cell-culture models. However, there remain several concerns of flow perfusion control, organ functions, and inter-tissue interaction that need to be addressed for GLA-MPS. First, gut and liver cells have different physiologically relevant flow perfusion to be realized via a precise flow control in the chip. Second, gut and liver cells have the cellular polarity, which should be functionalized in molecular transportation. Third, circulation flow needs to connect two organs for inter-tissue interactions. The current GLA-MPSs are based on two approaches: the cell-culture insert in the microtiter plates^7,11^ and microfluidic devices.^12,13^ For the cell-culture insert system, cellular uptake and transport across the cell barrier (e.g., endothelial cells and intestinal epithelial cells) can be studied with the porous membrane; under porous membrane, a circulation channel could be used to connect two organs as well as apply flow perfusion; however, it is difficult to apply an appropriate uniform perfusion flow for surface of each organ within the insert precisely, because of its macroscopic scale in the millimeter range and lack of perfusion controlling system. Practically gut epithelial cells require physiological relevant fluid shear stress (FSS) from the perfusion flow, to express their organ cell functions.^14,15^ In contrast, the use of a microfluidic device allows the control of perfusion flow in the planar compartmentalized chambers and the interconnection of two or more tissue cells via a microfluidic channel with an on-chip micropump.^12,13,16^ These microfluidic devices have some issues to be addressed; for example, we can apply only one flow rate for the introduced tissues although the gut and liver require different flow rates to realize their proper functions.^17,18^ Further, the device cannot be applied to study organ molecular transport function across the gut barrier because it does not have an apical to basal side transport structure such as a porous membrane substrate on the bottom channel.^12^ Thus, there is an urgent need to fulfill these requirements for establishing an advanced *in vitro* GLA model that (i) enables the application of appropriate perfusion flow for the gut and liver, (ii) studies molecular transports across the gut barrier, and (iii) interconnects the gut and liver to study their interactions within a single microfluidic device.

Here, we present a GLA-MPS device which can achieve individual physiological flow perfusion for each organ in the cell-culture chamber, form barrier tissue structure on a porous membrane, and connect two organs with circulation flow. The device is fabricated with four-layered microfluidic structures such as a cell-culture layer, porous membrane, circulation layer, and control layer with an on-chip micropump structure. The FSS on cells is estimated by the computational fluid dynamic simulation. The Caco-2 intestinal epithelial and HepG2 hepatocyte-like cells are applied to their own physiologically relevant FSSs for enhancing their functions, respectively. The circulation flow controlled by micropump performed closed-loop circulation to interconnect the Caco-2 and HepG2 cell-culture chambers via a porous membrane. This device allows modeling the GLA interaction as an *in vitro* model.

## Materials and Methods

### Device concept and structure

The GLA-MPS device was designed as shown in Figure 1a and b. The GLA-MPS device comprises four layers made of polydimethylsiloxane (PDMS) and a glass substrate (Figure 1c and Supplementary Fig. S1). The four PDMS layers work as a cell culture layer, porous membrane, circulation layer, and control layer integrated with a micropump. The PDMS meets the requirements as a structural material owing to its biocompatibility, gas permeability, and elasticity to realize micropumping.^19^ The cell-culture layer contains cell-culture chambers (length: 5.3 mm, width: 2 mm, height: 360 μm). A porous membrane has holes with a hole size of 5 μm, pitch of 20 μm, and thickness of 20 μm to prevent cell penetration through the holes and reduce the FSS for cells from the basal side. This membrane separates the cell-culture chamber and circulation channel to perform two different flows; soluble factors are transported via holes. An on-chip micropump was integrated into the control layer to perform closed-loop circulation. Further, the external pump was connected to the inlet of the gut chamber to apply the FSS on the apical side of the gut epithelial cells.

**Figure 1.**
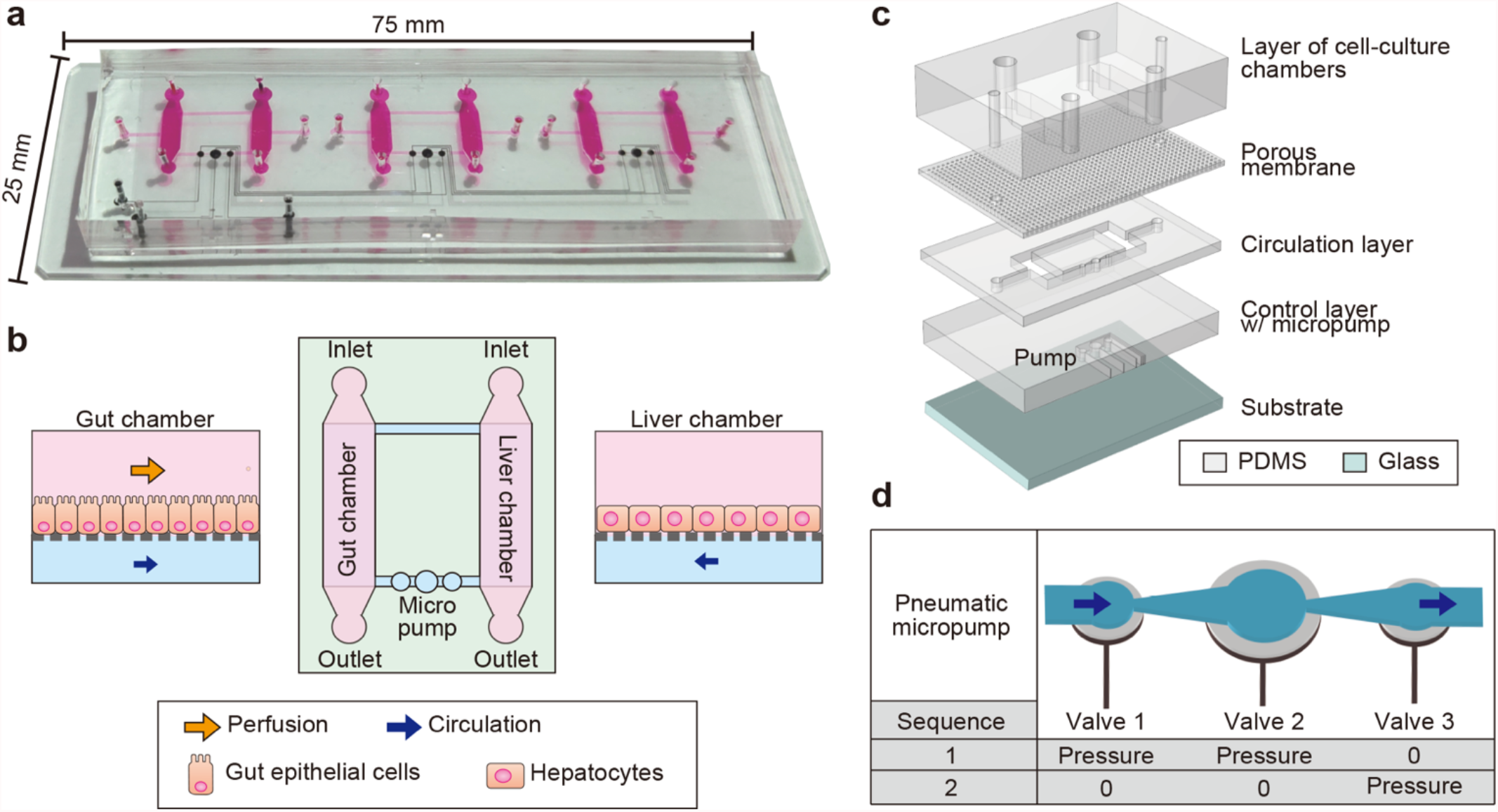
Conceptual illustration of the gut-liver-axis (GLA)-MPS. a. Photograph of the GLA-MPS with three sets of in vitro GLA and an integrated micropump. b. Illustration of microfluidic flow channels for establishing GLA-MPS. Gut epithelial cells and liver hepatocytes are cultured in corresponding cell-culture chambers, applied optimal perfusion flow for each cell type, and interconnected by circulation flow. c. Illustration of the GLA-MPS that comprises four PDMS layers and the bottom glass layer. d. Schematic of actuating micropump.

### Micropump actuation for closed-loop circulation

An approach that uses a pneumatic micropump enabled integration with a microfluidic device. We fabricated a pneumatic micropump, which is a modified version of ‘Quake’s microvalve pump’^20^ to perform closed-loop circulation (Figure 1d). The microvalve is integrated in the control layer, consisting of a flexible PDMS membrane and underlying control channel. Membrane actuation is achieved by varying the positive hydraulic pressure from the control channel, bending towards the circulation channel. This micropump uses three adjacent microvalves of the membranes in combination with a larger size (diameter: 1 mm) and two smaller ones (diameter: 0.6 mm), connected to a diffuser and nozzle structure^16^ for increasing the one-directional flow by reducing backward flow. The purpose of this ‘two phase’ actuating method is to simplify the actuating sequence, which reduces the efforts of frequency and pressure control from the pneumatic system. Here, circulatory perfusion flow is controlled by selectively releasing (“0”) and actuating (“Pressure”) microvalves in the membrane “push-up” configuration. “sequence-1” represents actuating both valves 1 and 2 simultaneously, which keeps valve 3 free. Then, “sequence-2” represents the state where only valve 3 is actuating. The actuation time of the 1-mm microvalve is slower than that of the 0.6-mm microvalves, because the larger volume of control layer chamber leads to a high inertial mass.^21^ Thus, valve 2 is slower than that of valves 1 and 3, which leads to a peristaltic valve motion to generate output flow in one-direction during actuation.

### Chip fabrication

The GLA-MPS device was fabricated using multilayer soft-lithography (Figure 2).^22^ For the resist mold of the cell-culture layer, a dry-film type negative photoresist (45 μm; TMMF S2045™, Tokyo Ohka Kogyo) was laminated 8 times to achieve a thickness of 360 μm; similarly, the TMMF was laminated 2 times to achieve a thickness of 90 μm for fabricating the resist mold of the circulation and control layers. The resist mold of the porous membrane layer was fabricated using a spin-coated 25-μm-thick negative photoresist (TMMR S2000™, Tokyo Ohka Kogyo). UV exposure was performed using a mask aligner (MUM-1000 Series, Japan Science Engineering)^23,24^ with Cr-patterned glass masks for the corresponding molds of each layer (cell culture layer: 1000 mJ cm^−2^, porous membrane: 1500 mJ cm^−2^, and circulation layer and control layers: 500 mJ cm^−2^). For the development, the resist molds were put in a propylene glycol monomethyl ether acetate (PGMEA) solution (PM Thinner, Tokyo Ohka Kogyo)) at 23 °C for 20 min. ^23,24^ The PDMS base and curing agent (Sylgard 184, Dow Corning) mixed in a ratio of 5:1 (w/w) for the porous PDMS membrane and 10:1 for other layers were poured onto resist molds and cured at 25 °C for 48 h. The designed membrane thicknesses were tuned by spin coating to 25 μm for the porous PDMS membrane and 60 μm for the pump membrane. The cured PDMS layer of the porous membrane was etched with a 5-μm thickness using O_2_ plasma (power 100 W: pressure 50 Pa: flow rate 50 sccm; FA-1, SAMCO) to open the through holes (Supplementary Fig. S2). The cell-culture layer was bonded to the porous PDMS membrane whereas the control layer was bonded to the circulation layer. The two assembled structures were peeled off from the substrate and bonded together using VUV surface activation (Min-Excimer SUS713, Ushio). Finally, the bonded PDMS layers were bonded on the glass substrate and cured in an oven at 45 °C for 24 h. Low-temperature bonding causes less deformations on the PDMS block, which prevents shrinkage and bending, and leads to flat surfaces for multilayer bonding.

**Figure 2.**
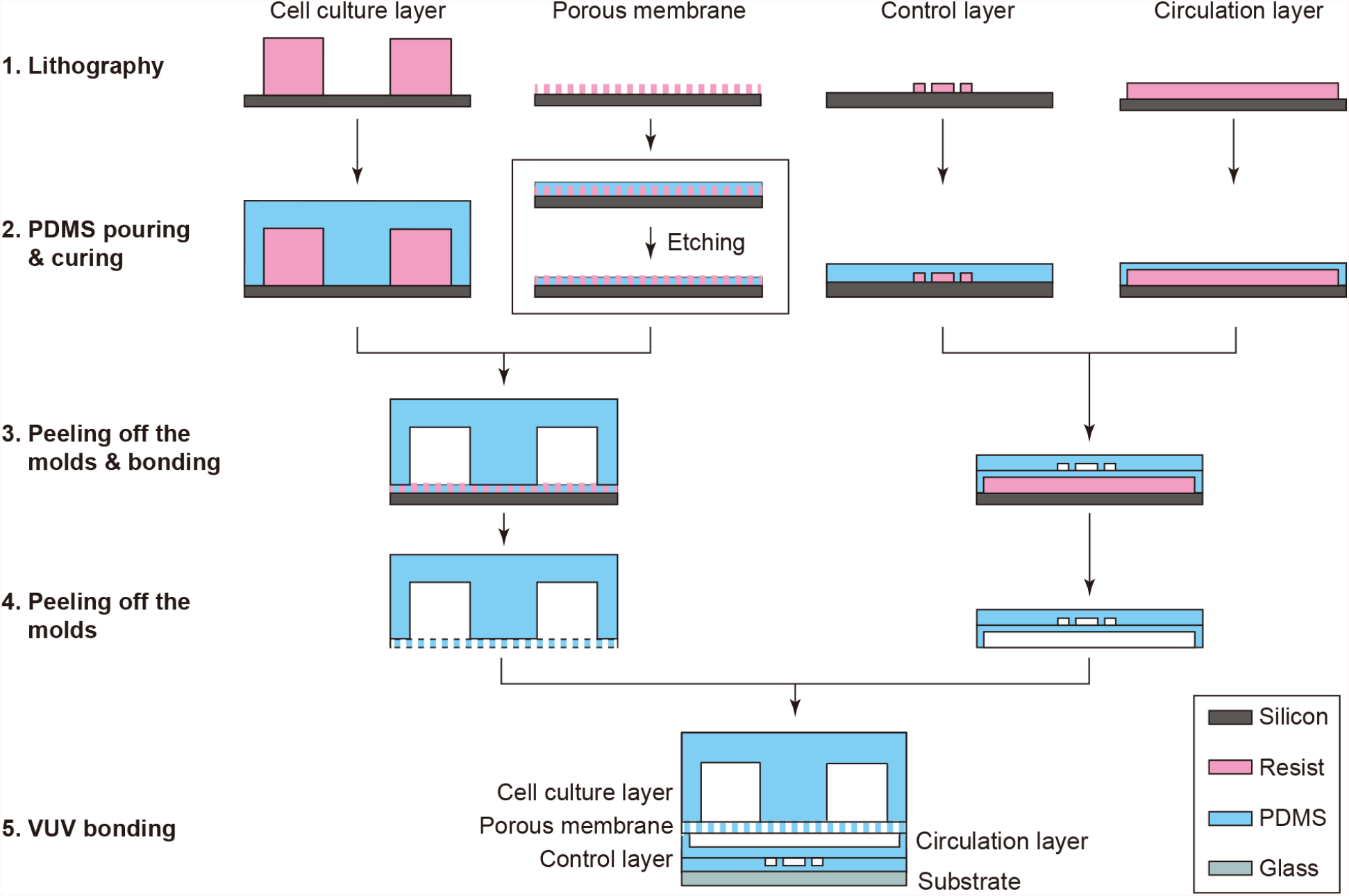
Fabrication procedure of the GLA-MPS. (1) Each mold was fabricated with photolithography. (2) Pre-PDMS mixture was poured on the molds and cured. (3) Solidified PDMS structures of the cell culture layer and control layer were peeled off from the molds and bonded on the porous membrane layer and circulation layer, respectively. (4) The bonded layers were peeled off from the mold, and then, (5) the layers were assembled on the glass substrate using VUV bonding.

### Evaluation of micropump flow rate

Microbead tracing (4.5 μm in diameter) was used for visualizing the culture medium flow to characterize the flow rate generated by the on-chip micropump; the flow distances of the beads were measured. Before the experiments, the microfluidic channel was coated with 1% (w/v) bovine serum albumin (BSA, Sigma-Aldrich) in PBS for 2 h at 25 °C to prevent the nonspecific adhesion of beads on flow channels and cell-culture chambers. The beads were suspended in 1% (w/v) BSA solution at 1 × 10^6^ beads mL^−1^; subsequently, a bead-suspended solution was introduced into the device. The movement of the microbeads was captured using a CMOS camera (EO-4010M, Edomond Optics) equipped with a microscope (GX53, Olympus); the bead velocities for flow velocity were analyzed using a custom-made MATLAB program (R2020b, MathWorks). The volumetric flow rates were calculated by multiplying the flow velocity by the cross-sectional area of the microfluidic channel.

### Computational fluid dynamics for fluid shear stress

Computational fluid dynamics (CFD) analysis was used to estimate the FSS of the cell surface using COMSOL Multiphysics (Ver. 5.6, COMSOL Inc.). The center cross-section along the gut chamber perfusion direction was used in the calculation; the microhole structure of the porous membrane was incorporated. The cell-culture medium was considered an incompressible and homogeneous Newtonian fluid with conditions similar to water at 37 °C (density: 997 kg m^−3^, viscosity: 6.9 × 10^−4^ Pa s^−1^). A 10-µm-thick layer covered the upper side of the porous membrane with no direct flow across via the interior wall setting to model the cell layer. No-slip boundary conditions were applied to the channel walls. A laminar flow/stationary library was used to determine the shear rate 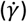, and the FSS (τ) was calculated by multiplying the shear rate with the dynamic viscosity of water.

### Cell culture

HepG2 human hepatocellular carcinoma and Caco-2 human colorectal adenocarcinoma cell lines were obtained from the American type culture collection (ATCC). Cells were maintained in the cell culture medium containing Dulbecco’s modified Eagle medium (DMEM) (Sigma-Aldrich, St. Louis, MO, USA) supplemented with 10% (v/v) fetal bovine serum (FBS, Cell Culture Bioscience), 1% (v/v) nonessential amino acids (NEAA, Thermo Fisher Scientific), and 1% (v/v) penicillin/streptomycin (P/S, Thermo Fisher Scientific) in a humidified incubator at 37 °C with 5% (v/v) CO_2_. HepG2 and Caco-2 cells were passaged with trypsin/EDTA (0.04%/0.03%[v/v]) solution every 3 and 7 days, respectively.

### Cell culture on a chip

The chips were sterilized with 70% (v/v) ethanol and an ultraviolet light in a biosafety cabinet at 25 °C for 40 min prior to the cell culture on the chips. Then, two chambers in the cell-culture-chamber layer were filled with 1.3% (v/v) Matrigel hESC-qualified matrix (Corning) in DMEM/F12 (Sigma-Aldrich) at 25 °C for 2 h. The cell culture and circulation chambers were washed and filled with a cell-culture medium. The chips were incubated at 4 °C for 16 h to remove all air bubbles from the chips. The next day, after washing the chips with a cell-culture medium warmed at 37 °C, three microliters of cell suspensions of Caco-2 and HepG2 cells (5 × 10^6^ cells mL^−1^) were introduced into the corresponding cell culture chamber chips; the cell density was 1800 cells mm^−2^. The chips were placed in a humidified incubator at 37 °C with 5% (v/v) CO_2_. After 24 h, the cell culture medium was changed to remove the floating dead cells. For the circulation and perfusion, the inlet of the gut culture chambers was connected to a syringe pump (KDS LEGATO 210, KD Scientific Inc.); the micropump was actuated for medium recirculation. For a static cell culture, the cell culture and circulation chamber layers were changed with the cell culture medium every 6 h. The total cell culture on the chip was maintained for up to 10 days.

### Live/dead cell assay

A cell viability assay was conducted using the NucleoCounter NC-200, Via1-Cassette™ (Chemometec). After the cell culture or treatment, cells in the chips were washed with PBS once, by adding 50 μL PBS through the cell culture chamber and the circulation channel. Then, 20 μL of trypsin/EDTA solution was added through the cell culture chambers and incubated at 37 °C for 10 min. After cell detachment, cells in both chambers were separately harvested into a 1-mL tube and diluted to 200 μL with the cell-culture medium. The homogenous cell suspension was aspirated into a Via1-Cassette™ to count the number and ratio of dead/live cells.

### Immunocytochemistry

After the cell culture and treatment, the cell-culture chambers and circulation layer were washed twice with PBS containing 50 µL of PBS. The cells were fixed with 4% paraformaldehyde (PFA, FUJIFILM Wako Pure Chemical) in PBS for 15 min at 25 °C and permeabilized with 0.1% (v/v) Triton X-100 in PBS for 30 min at 25 °C. The cells were then incubated in a blocking buffer (5% normal goat serum, Vector; 5% normal donkey serum, Wako; 3% BSA, Sigma-Aldrich; and 0.1% Tween-20, Nacalai Tesque, Inc., in PBS) at 4 °C for 16 h. After blocking, the cells were incubated with primary antibodies such as rabbit anti-human ZO-1 IgG (Thermo Fisher Scientific) and mouse anti-human albumin IgG (Thermo Fisher Scientific) in a blocking buffer at 4 °C for 16 h. After washing the primary antibodies with PBS-T twice, the cells were incubated with secondary antibodies such as Alexa Fluor 488-labeled donkey anti-rabbit IgG and Alexa Fluor 647-labeled donkey anti-mouse IgG H&L (Jackson ImmunoResearch) in a blocking buffer at 25 °C for 1 h. The cell nuclei were stained with 300 nM 4,6-diamidino-2-phenylindole (DAPI, Dojindo Laboratories). The DAPI solution (50 µL) was introduced into the chambers and incubated at 25 °C for 30 min; the chambers were washed with PBS.

### Imaging acquisition

Confocal microscopic imaging was performed using an Andor Dragonfly 502 system (Andor, UK). This system is built on a Nikon ECLIPSE Ti2-E inverted fluorescence microscope (Nikon, Tokyo, Japan) and equipped with a Zyla 4.2 Plus scientific complementary metal oxide semiconductor (sCMOS) camera (Andor) driven by Fusion (Andor), a CFI Plan apochromat mambda 10x dry/0.45 NA (Nikon), motorized XY stage and stage piezo (Applied Scientific Instrumentation, MO, USA), and three lasers (405, 488, and 637 nm). For image acquisition, the lasers were set to 488 nm for ZO-1, 637 nm for albumin, and 405 nm for DAPI.

### Permeability assay across cell barrier

FITC-dextran (4 kDa, Sigma-Aldrich) was used for the permeability assay.^25^ It was dissolved into a cell-culture medium at a concentration of 1 mg mL^−1^ as the working solution. The working solution was introduced into the gut cell-culture chamber and incubated for 4 h. The FITC-dextran in the medium of the cell-culture chamber and circulation channel was separately collected and measured using the Synergy HTX microplate reader (BioTek Instruments). The hydraulic conductivity *K*_*dextran*_ was calculated using

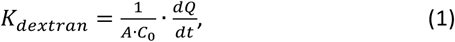

where *A, C*_*0*_, and *dQ/dt* represent the transporting surface area of the porous membrane (5.3 mm × 2 mm), original concentration of the FITC-dextran in the gut apical chamber, and amount of FITC-dextran transported across the membrane over the respective time interval, respectively.

### Statistical analysis

The Tukey–Kramer test and asymptotic Wilcoxon signed-rank test were conducted using R software (ver. 3.5.2; https://www.r-project.org/).

## Results & Discussions

### Computational estimation of shear stress and flow rates in GLA-MPS

The FSS was one of the most important physical factors affecting the cell function and morphology,^26,18^ but the previous GLA-MPSs faced challenges in the control. The flow perfusion conditions should be investigated for each organ cells. In this device, we estimated the FSS and flow rates using COMSOL simulation to investigate whether the GLA-MPS provides proper FSS on cells (Figure 3). When the medium perfusion in the gut chamber was performed from 2 to 5 μL min^−1^, the surface FSS on the apical side of cells had a linear relationship with the external pumping flow rate (Figure 3a and 3b). The mean FSS for the intestinal cells was in the range of 8 × 10^−3^ to 2 × 10^−2^ dyne cm^−2^, which was in the physiologically relevant range^26,27^ and could promote gut cell barrier formation and differentiation process.^28,29^

**Figure 3.**
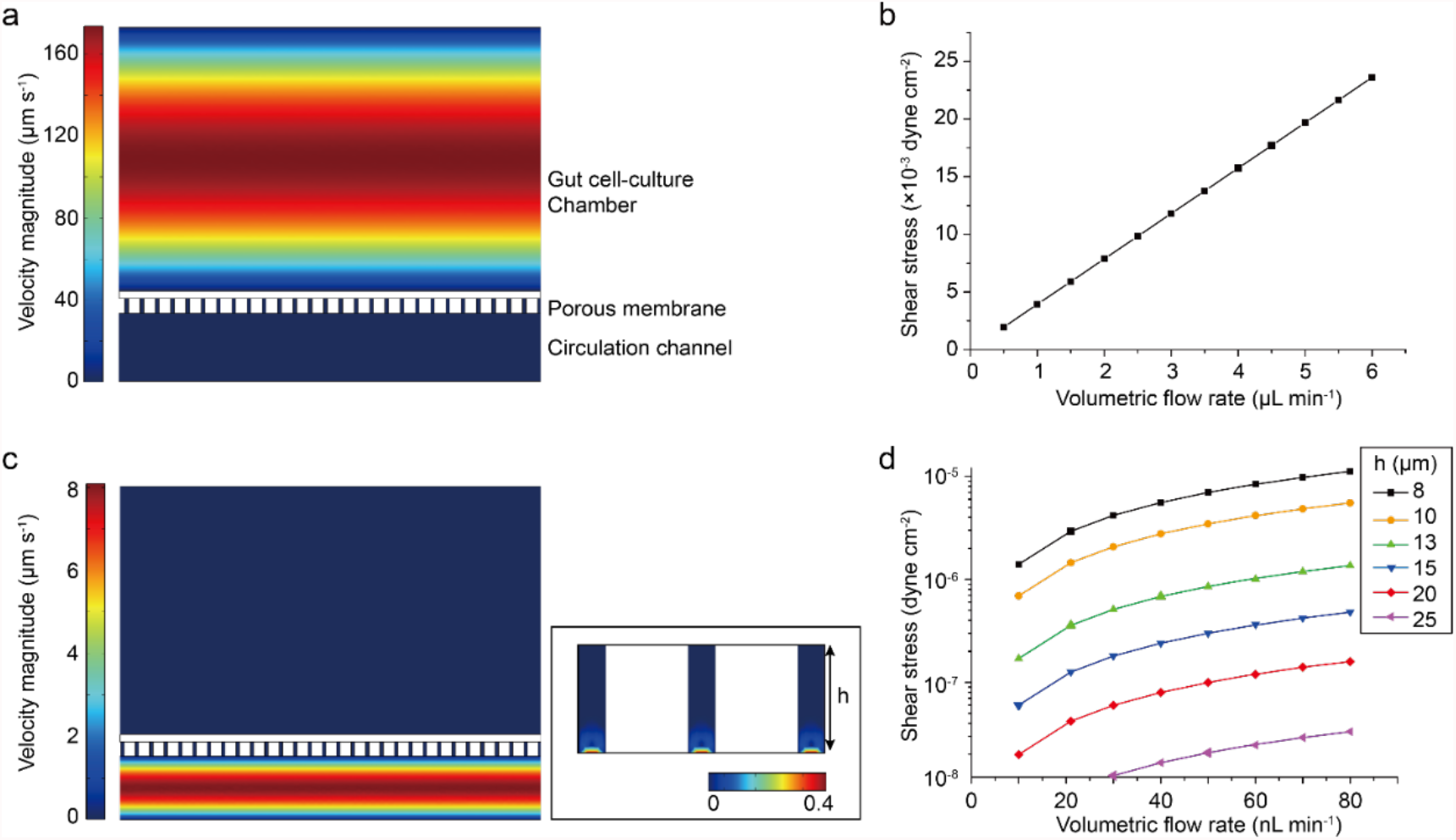
Simulated perfusion flow in the GLA-MPS. (a) Simulated perfusion flow in the gut cell culture chamber. (b) Simulated shear stress on the cell surface cultured on the porous membrane. (c) Simulated circulation flow in the circulation chamber. (d) Simulated shear stress by the circulation flow.

The FSS applied to cells from the basal side through the holes of the membrane was determined using the membrane thickness (h) and circulation flow rate (Figure 3c and d). The FSS was reduced dramatically with an increase in membrane thickness. In our setup parameters (20-µm-thick membrane), we found that the surface FSS (1 ×10^−8^ to 1.2 × 10^−7^ dyne cm^−2^) on the basal sides of cells by the circulation flow were considerably smaller than those of the FSS on the apical side of gut epithelial cells. Thus, in the effect of circulation flow, we not only considered the effects of FSSs of basal sides for the later cell experiments, but we also recognize that circulation flow would mediate communication between the gut and liver cells.

In recent results for the gut on a chip^17^, the Caco-2 cells on the porous membrane experienced the flow perfusion in the apical side or basal side, which showed an improvement of cell functions. However, so far, there are seldom reports of GLA-MPSs used to investigate the flow perfusion for cells on the apical side or flow on the basal side. In our device, by using the simulation model, the FSS on cells is well-designed and can be clearly used for investigating their effect on cells from both sides.

### Evaluation of circulation flow by the micropump

The micropump for circulation flow was controlled using the developed pneumatic actuating method to induce the flow circulation between two organ chambers. We investigated the circulation flow rates controlled by an on-chip micropump after the fabrication of the GLA-MPS (Figure 4). The mean flow rate increased with driving pressure (0–150 kPa) and pumping frequency (0–1.5 Hz). Working frequencies over 1 Hz exhibited no significant differences in the generated flow rates; in contrast, the GLA-MPS had disassembled by applying over 150 kPa in the control layer for 14 days (data not shown). Therefore, the working pressure was set from 50 to 150 kPa to carry out cell culture and assay in the GLA-MPS; this gave flow rates from 22 to 61 nL min^−1^ for circulation at a 1-Hz working frequency.

**Figure 4.**
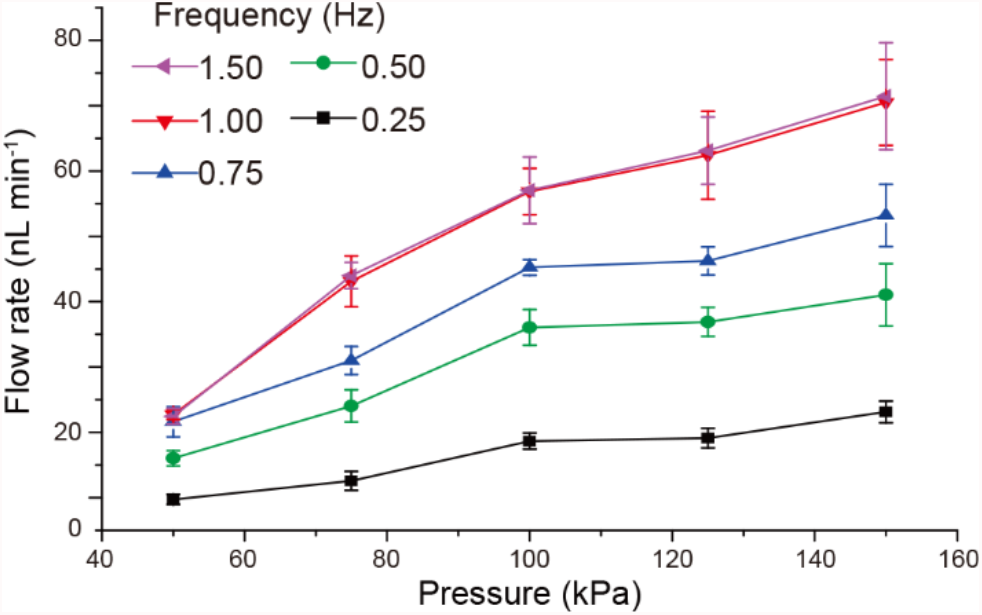
Measured mean flow rate of the micropump for pressure-dependent actuation from 0.25 to 1.5 Hz frequency. Each dot represents the mean ± standard deviation (SD) (n = 10).

The flow rate requirement for inducing two organ interactions could be estimated from the exchange ratio of circulation flow rate (q) and flow volume (Q), i.e. q/Q. In this device, the microfluidic system has a circulation flow volume of 1.5 µL. The exchange ratio is 1.5% to 9% in 1 min. In other GLA-MPSs,^3,7^ the exchange ratio is approximately 0.25% to 1% in 1 min. Thus, our device realized a more intensified circulation flow between the gut and liver. For 9-day cell culture, this flow is sufficient to induce the interaction between two organs in the GLA-MPS device. Besides, the on-chip micropump could realize a fine range of flow rate adjustment, which is consistent with the aforementioned FSS simulation circulation flow rate range. Therefore, we could couple the circulation flow with gut perfusion flow to improve the cell function and induce the organ to organ interaction for cell culture experiments.

### GLA culture on a chip

Caco-2 gut intestinal^30^ and HepG2 hepatocyte-like cells^31^ were used and cultured in GLA-MPS to establish *in vitro* GLA. Prior to the cell culture, the surface of the porous membrane should be coated with Matrigel for cell attachment. By applying the pressure barriers of caplillary^32^, the Matrigel was only coated on the surface of the cell-culture chamber side (upper side), but not the surface of holes and circulation channel side (bottom side). This is for reducing cell attaching and migrating through the hole of the porous membrane (Supplementary Fig. S3).

After culturing for 1 d under the static conditions for the cell attachment, only Caco-2 cells, but not HepG2 cells, were subjected to the perfusion flow (Figure 5a). Both Caco-2 and HepG2 cells were maintained for eight days with the circulation flow in the circulation channel. After the culture, Caco-2 and HepG2 cells showed 95% and 92% cell viability, respectively (Figure 5b). It demonstrated that the cells could be maintained relatively high cell viability in this device.

**Figure 5.**
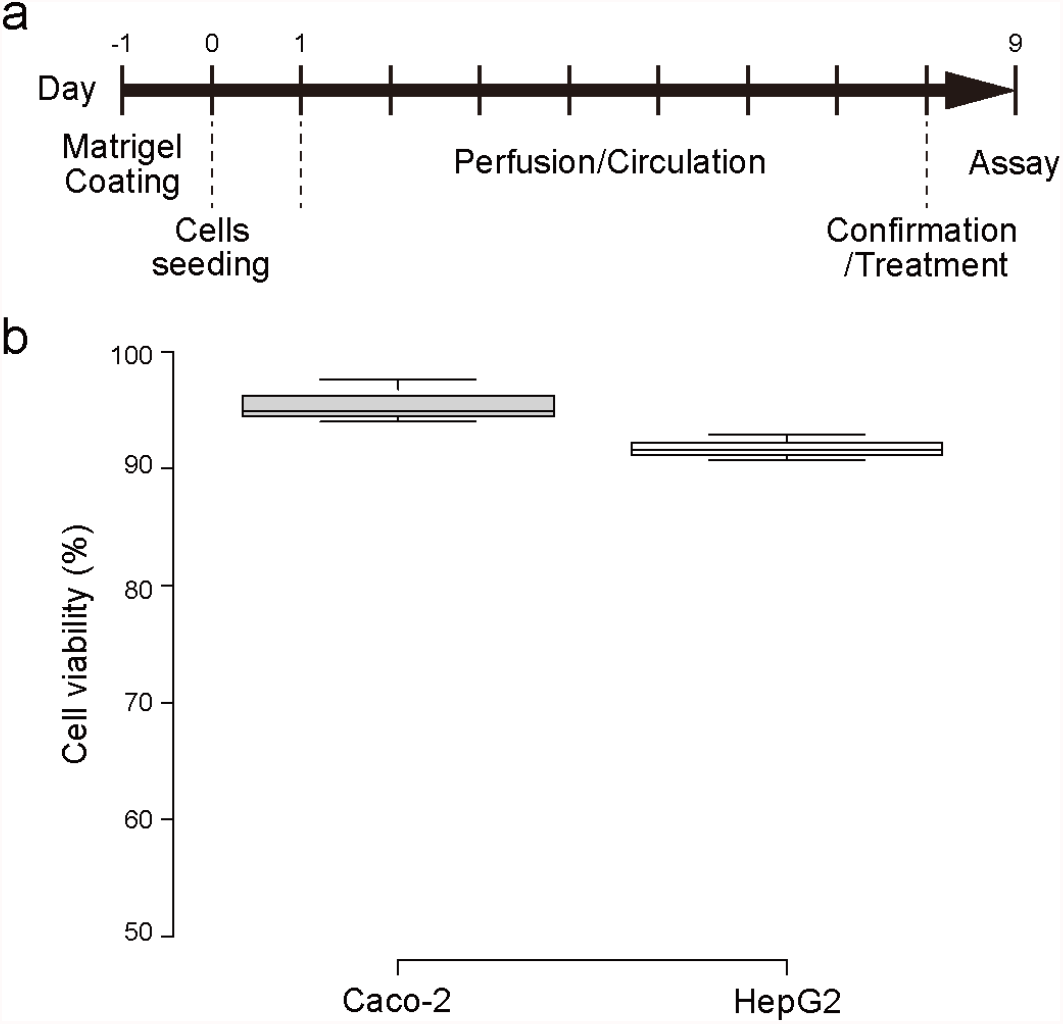
(a) Experimental procedure of cell culture, perfusion, circulation and treatments in the GLA-MPS. (b) Evaluation of cell viability of Caco-2 and HepG2 cells in the GLA-MPS at 2 μL min^−1^ perfusion and 61 nL min^−1^ circulation flows.

The expression of ZO-1 and albumin (ALB), as functional markers of the gut and liver, respectively, were observed by immunocytochemistry to confirm the functions of cells cultured in the GLA-MPS (Figure 6). Caco-2 cells expressed ZO-1 tight-junction protein to form the gut barrier (Figure 6a); the increase in flow rates of perfusion flows in the gut cell culture chamber induced the ZO-1 expression in the Caco-2 cells at 5 μL min^−1^ (FSS = 2 × 10^−2^ dyne cm^−2^). Further, the circulation flow did not change the ZO-1 expression in the Caco-2 cells without perfusion flow; instead, it induced ZO-1 expression in addition to the perfusion flow. In light of the simulation results, FSS by circulation flow was considerably smaller than that by perfusion flows; however, the cultured cells could receive the FSS of basal sides. The cell-culture medium in the circulation channel could accumulate molecules (i.e., growth factors, cytokines, metabolites, and lipids) and exosomes secreted from Caco-2 and HepG2 cells. Thus, the medium contained both autocrine and paracrine factors and facilitated the interactions of Caco-2 and HepG2 cells by the circulation flow. Thus, the induction of ZO-1 expression could be the outcome of the combinations of FSS by perfusion/circulation flows and molecular transport.

**Figure 6.**
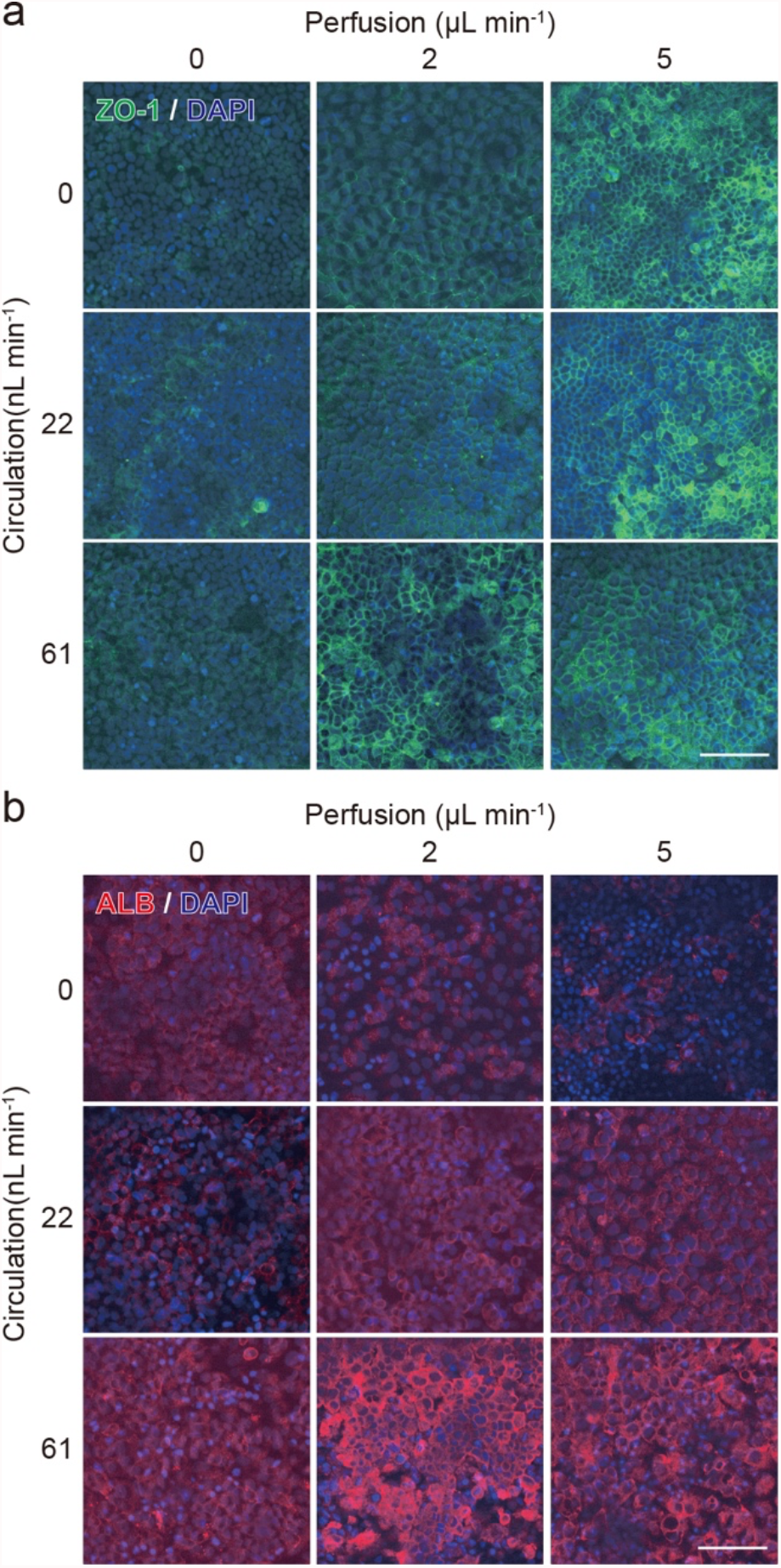
Optimal perfusion and circulation flows enhanced the expression of ZO-1 (Caco-2) and albumin (ALB; HepG2). Scale bar 100 µm.

In the case of HepG2 cells, the HepG2 cells were not directly subjected to perfusion flow in the cell-culture chamber; HepG2 cells did not change the ALB expression by changing the perfusion flow rates (Figure 6b). In contrast, the increase in circulation flow rates from 0 to 61 nL min−^1^ induced ALB expression in HepG2 cells. These results suggest that HepG2 cells facilitated hepatic functions such as ALB production by FSS, and as a molecular transport in the circulation channel. The conventional cell-culture methods continue to have an important but unsolved challenge to activate *in vitro* cultured hepatocytes to express their *in vivo* physiological hepatic functions. Our approach using chemical and physical stimuli suggested a new way to obtain hepatic functions *in vitro*. The conventional MPSs could not provide these results because of the aforementioned drawbacks of MPSs; our MPS showed advantages over the conventional MPSs.

We select the flow rates (Gut perfusion flow: 2 µL min^−1^; circulation flow: 61 nL min^−1^) to establish *in vitro* GLA, which provides the highest expression of functional proteins in both Caco-2 and HepG2 cells among the tested conditions. Using this optimal condition, we tested the permeability of Caco-2 cells after the 9-day culture, and we confirmed the cell barrier function (Figure 7). It indicated that Caco-2 cells formed an intact cell barrier to protect the transportation of 4-kD FITC-dextran across the porous membrane.

**Figure 7.**
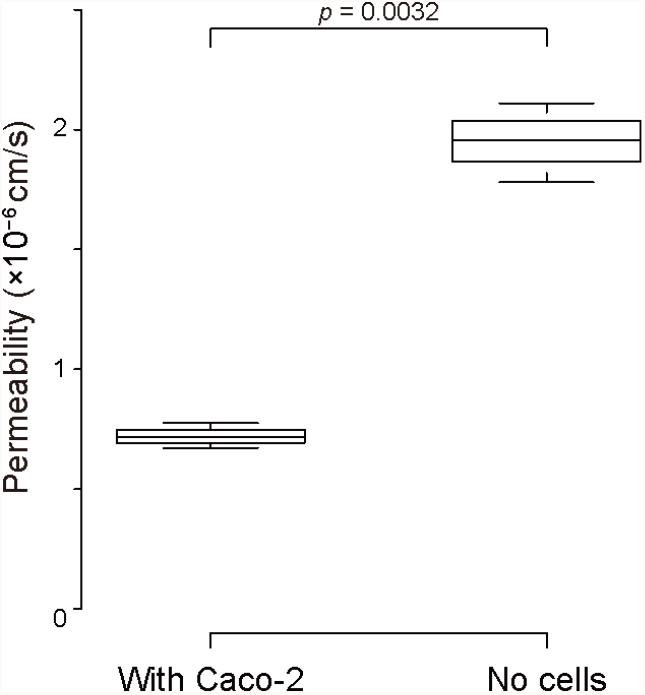
Permeability assay to evaluate Caco-2 barrier function by observing FITC-dextran transportation from the culture chambers to the circulation channel via the porous membrane. n = 3.

Thus, both Caco-2 and HepG2 expressed better functions than those in the conventional cell culture settings by applying optimal flows and inter-tissue interactions to establish *in vitro* GLA. Further, this feature is advantageous for modelling multiple organ diseases such as inflammatory bowel disease (IBD) and non-alcoholic fatty liver disease (NAFLD). We can treat one of the tissues to induce disease phenotypes and investigate its effects on other tissues. Gut microbiome^33^ plays critical roles in the pathophysiological scenarios in the gut; they can influence other organs such as the liver^34,35^ and the immune system^37^. Although it is well known that IBD^7^ and NAFLD^35^ are strongly associated with the gut microbiome, effective drugs or treatments are not available in the market because of the lack of mechanistic insights. Since the presented GLA-MPS has an individual addressability for cell-culture chambers, we will introduce a microbiome in only the gut cell-culture chambers and investigate the disease mechanisms in GLA altered by microbiota. Instead of liver cells; our MPS device allows the introduction of other tissue cells such as neurons, and we can investigate the interactions with the gut microbiome. Thus, our MPS is not limited to recapitulate only GLA, but it is also applicable to other organs and the microbiome.

Further, the presented GLA-MPS allows a collection of conditioned cell-culture media from two cell-culture chambers and circulation channels, separately, for investigating the underlying molecular mechanisms of the GLA by biochemical assays such as ELISA and mass spectroscopy. Since these methods for the cell-culture medium are non-invasive for cultured cells, we will be able to determine the transitions of molecular contents in the medium during the experimental period to explore interesting molecular biomarkers. At the end of the experiments, GLA-MPS allows performing invasive cell-analytical methods such as immunocytochemistry, as shown in this paper, as well as multi-omics analysis (i.e., genomics, transcriptomics, proteomics, and metabolomics) to obtain deeper insights into normal and diseased GLA.

One advantage of using microfluidic technology for disease modelling is the minimization of the number of cells, such as the primary cells obtained from the patients. These cells are limited in their ability to recapitulate the patients’ pathological conditions *in vitro*. The presented MPS requires only 1.5 × 10^4^ cells for each cell culture chamber, which is considerably smaller than those of the 24-well cell culture inserts (3 × 10^5^ cells). Therefore, we efficiently use precious patient samples, such as biopsy specimens. In the present study, we used widely used cell lines (Caco-2 and HepG2) that can provide the standard performance of the presented GLA-MPS. Since patient-derived primary cells could have large patient-to-patient variations, it would be beneficial to obtain such standard datasets with the use of commonly used cell lines.

The device is fabricated using the microfabrication method with a four-layered planer microfluidic structure. Owing to this feature, its another advantage is the capability to integrate electrical read-out sensors in cell-culture chambers and circulation channel, which allows monitoring cell conditions and behaviors during cell culture and treatments in a real-time and non-invasive fashion.^38^ Such sensors will provide temporal information on tissue formation, disease progression, and drug treatments. In the case of the GLA, the sensor for measuring trans-epithelial electrical resistance (TEER) was beneficial for evaluating gut barrier formation and causing damage during disease progression. In this study, we used a fluorescent dye to investigate gut barrier function; however, the dye might also influence cellular functions. Until now, gut-on-a-chips integrated with a TEER measurements have been reported.^39,40^ The designs and fabrication processes need to be further improved for better accuracy and reproducibility, and to integrate it into multi-organ MPS. This integrative approach will lead to the advancement of MPS for a better understanding of disease mechanisms and applications in drug discovery.

## Conclusions

We developed a GLA-MPS platform that can (i) cultivate gut and liver cells separately with applied different perfusion flows/FSS to improve their cell viabilities and functionalities, (ii) form barrier tissue, and (iii) interconnect GLA via circulation channels. The device is a more distinctive platform than the current GLA-MPS, resembling the physiological flow perfusion and circulation of *in vivo* GLA. It is also suitable for demonstrating the individual accessibility to investigate the GLA interaction, mediated by IBD and NAFLD. However, the GLA-MPS still has some limitations that need to be improved in future. For example, the device can be integrated with the organoids structure, which could extend the gut and liver organ functions for tissue engineering and disease modeling. In addition, the device should have capabilities beyond FSS control and should also control the molecular transport gradient across the barrier tissues effectively in the microenvironment to understand the GLA interaction. Finally, our approach based on GLA-MPS has the potential for applications in understanding the GLA mechanisms associated with individual physiological flow perfusion and multi-organ interactions for disease modeling.

## Supporting information

Supplementary information for Gut-liver-axis microphysiological system for studying cellular fluidic shear stress and inter-tissue interaction

## Author Contributions

J.Y., Y.H., T.T., O.T., and K.K. conceptualized the study. J.Y. and Y.H. fabricated the devices. J.Y. and S.I. performed the cell culture experiments and analyses. J.Y. and K.K. performed the image analysis. All authors contributed to data analysis, discussion, and interpretation. J.Y., Y.H., and K.K. wrote and revised the manuscript with input from all authors.

## Conflicts of interest

There are no conflicts to declare.

## Acknowledgments

Funding was generously provided by the Japan Society for the Promotion of Science (17H02083, 18KK0306, 19H02572, and 21H01728), the Ebara Hatakeyama Memorial Foundation, Japan Agency for Medical Research and Development (17937667), and LiaoNing Revitalization Talents Program (XLYC1902061). Part of this work was supported by the Nanotechnology Platform Project within MEXT, Japan, through the Kyoto University Nano Technology Hub. The authors thank the iCeMS Analysis Center for access to the analytical instruments. WPI-iCeMS is supported by the World Premier International Research Centre Initiative (WPI), MEXT, Japan.

## Notes

### Competing Interest Statement

The authors have declared no competing interest.

