## Supplementary information for Gut-liver-axis microphysiological system for studying cellular fluidic shear stress and inter-tissue interaction for "Gut-liver-axis microphysiological system for studying cellular fluidic shear stress and inter-tissue interaction"

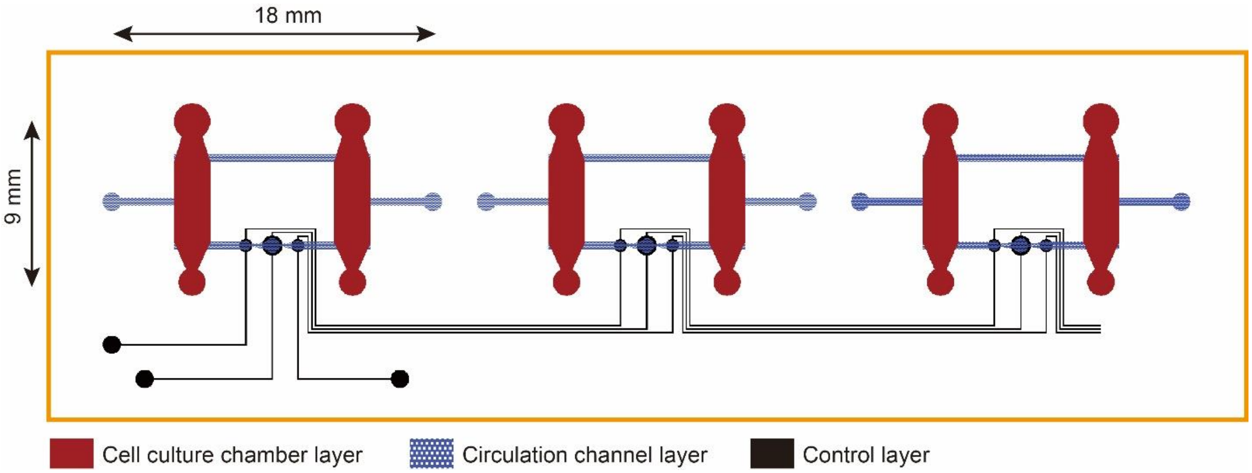


**Supplementary Fig. S1. Mask pattern design of GLA-MPS.**

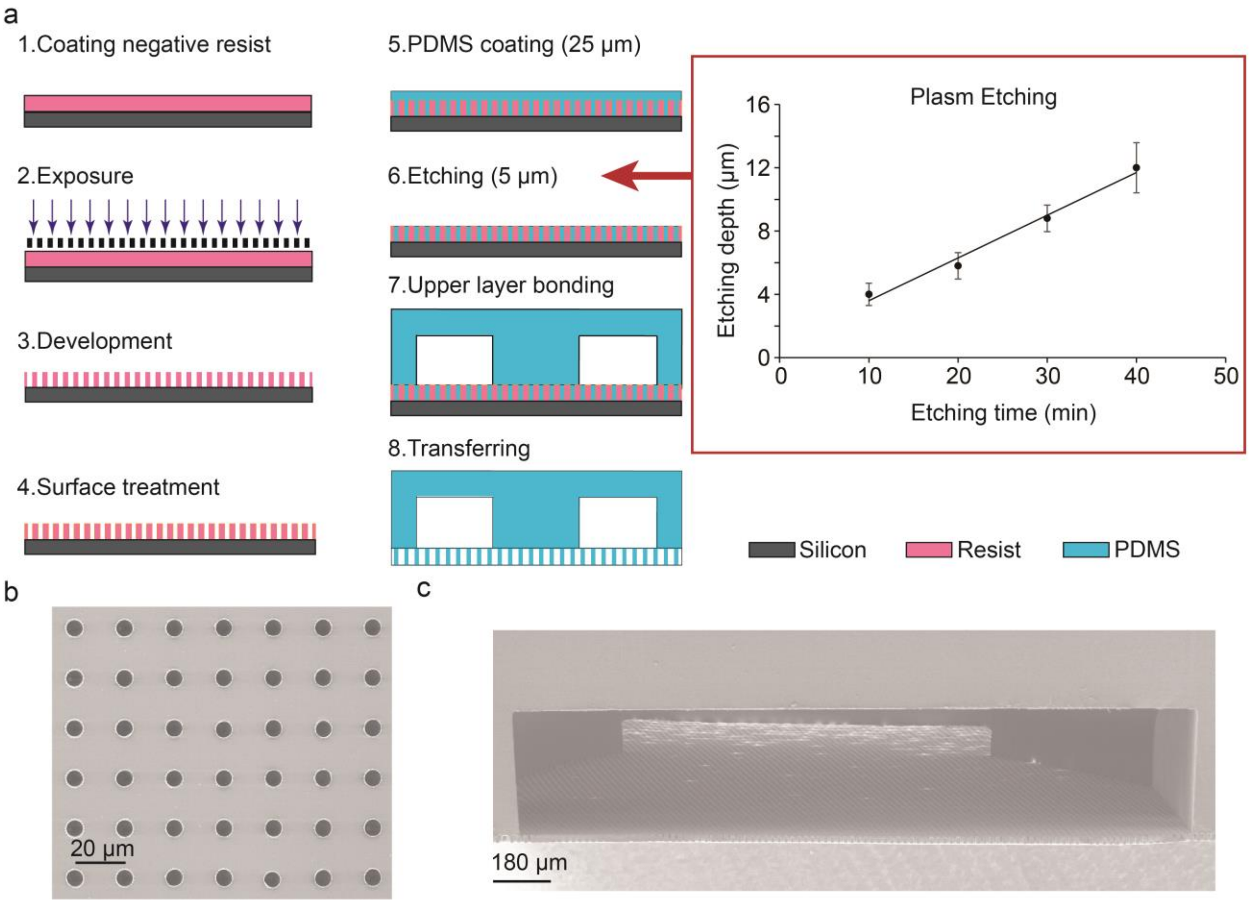


**Supplementary Fig. S2. Microfabrication of porous PDMS membrane. a.** (1) Negative photoresist was uniformly coated on the silicon wafer with a thickness of 24 ± 1 μm; (2) Contact exposure was performed to generate patterns under a photomask; (3) After post-baking and development, the resist mold was fabricated; (4) The surface of resist mold was treated with Shin-Etsu Barrier Coat No. 7 for easily peeling off the PDMS; (5) A 25-μm-thick PDMS layer was spin coated on the resist mold wafer; (6) Plasma etching

($O_2$: $CF_4$ = 1:1) with a flow rate of 50 ml/min was used to etch the PDMS 5 µm and open the through holes; (7) The cell culture layer was bonded on the porous membrane after VUV surface activation; (8) Finally, the assembled structure was transferred to the cell culture layer and peeled off from the wafer. **b.** Scanning electron micrographs (SEM) of the porous PDMS membrane after plasma etching. **c.** Cross-section view of assembled structure with cell culture chamber after transferring a porous membrane.

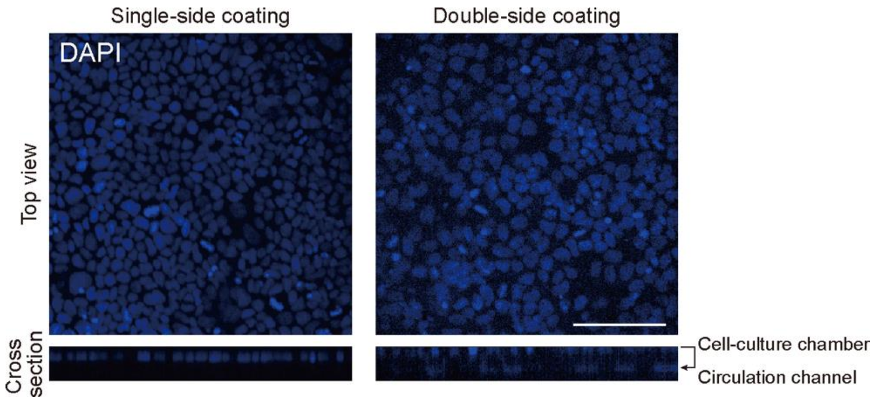


**Supplementary Fig. S3. Single-side coating of Matrigel on the porous membrane in cell culture chambers prevent cell migration across the pores of the porous membrane by applying pressure barriers of the capillary.** Cell nuclei staining shows that single-side coating prevented the Caco-2 cell from penetrating across the pore of the porous membrane compared with double-side coating. Scale bar: 100 µm.
